# Myeloperoxidase impairs mucociliary transport on human airway epithelium

**DOI:** 10.1101/2025.11.17.688903

**Authors:** Allison Boboltz, Vaidehi Rathi, Sahana Kumar, Gregg A. Duncan

**Affiliations:** Fischell Department of Bioengineering, University of Maryland, College Park, MD 20742; Biological Sciences Graduate Program, College Park, MD 20742

## Abstract

Dampening neutrophil-driven inflammation in the airways remains a challenge in treating cystic fibrosis (CF) lung disease. Myeloperoxidase (MPO) is a neutrophilic enzyme that produces reactive oxygen species and is highly concentrated in CF sputum samples. Greater MPO concentrations have been previously correlated with increased mucus plugging in bronchiectasis, suggesting that the enzyme could impair mucociliary transport. MPO reacts competitively with either thiocyanate (SCN^-^) or chloride (Cl^-^) in the airways to catalyze the production of hypothiocyanous acid (HOSCN) or hypochlorous acid (HOCl), respectively. HOCl has proved in prior studies to be extremely cytotoxic, while HOSCN can drastically reduce cytotoxicity. The concentration of SCN^-^ in the airways is largely dependent on transport by the cystic fibrosis transmembrane conductance regulator (CFTR) protein, which is dysfunctional in individuals with CF and causes low SCN^-^ concentrations. CFTR modulator therapies likely raise the concentration of SCN^-^ and enhance the production of HOSCN in the airways. We found that MPO inhibits mucociliary transport *in vitro* in regardless of SCN^-^ concentrations primarily due to increasing the macromolecular components and effective viscosity of airway surface liquid. The impairment of mucus clearance by MPO was similar to neutrophil elastase (NE), another neutrophilic granular enzyme that damages the host tissues and induces the secretion of mucin proteins by the airway epithelium. Overall, these findings identify MPO as a therapeutic target to resolve deficits in airway clearance function in CF and other related muco-obstructive lung diseases.

## Introduction

Cystic fibrosis (CF) is an incurable genetic disorder caused by malfunctioning of the cystic fibrosis transmembrane conductance regulator protein (CFTR).^1^ In the lungs, CFTR controls the ionic composition of the airway surface layer by transporting anions such as chloride and thiocyanate.^2^ Maintaining proper hydration and ionic gradients in the airway surface layer are important in supporting normal mucociliary transport (MCT).^3,4^ During MCT, mucus is cleared by airway epithelium via the coordinated beating of cilia in the periciliary layer. Mucus forms a protective gel barrier lining the airways to trap inhaled particles and microbes before being removed from the airways via MCT. Mucus consists of approximately 98% water and 2% solids in health.^4^ The solid components of mucus include various proteins such as mucin glycoproteins and DNA from dying airway epithelial and immune cells. In CF, CFTR dysfunction leads to highly viscoelastic mucus with an increased percentage of solid components (often >10% solids) that is not effectively cleared by MCT.^4^ Another downstream effect of the thick, stagnant mucus is uncontrolled inflammation in CF airways. Inhaled bacteria that become trapped in the dense mucus can colonize the airways and form biofilms that lead to chronic infections, causing inflammation. In addition, the viscous mucus layer can cause hypoxia-induced necrosis of the airway epithelial cells, leading to “sterile inflammation” with no infection.^5^ The excess release of inflammatory mediators in CF airways continues to promote the secretion of various proteins and release of DNA that further enhance the viscoelasticity of the mucus in a cyclic manner.

The inflammatory response in CF is dominated by neutrophils that migrate across the bronchial epithelium into the mucosal barrier. Neutrophil effector functions fall into three main subcategories: degranulation, neutrophil extracellular trap release (NETosis), or phagocytosis.^6^ All neutrophil effector functions involve the use of granules containing enzymes including myeloperoxidase (MPO), which is one of the most abundant proteins in neutrophils.^7^ Both degranulation and NETosis involve the extracellular release of MPO, causing elevated levels in CF airways.^8,9^ While degranulation releases free MPO, NETosis releases MPO that is complexed with DNA from de-condensed chromatin and greatly reduces its enzymatic activity.^10,11^ However, a large population of people with CF inhale DNase daily to degrade the chromatin structure of NETs, which releases active MPO.^11^

MPO reacts with the substrates hydrogen peroxide (H_2_O_2_) and halide or pseudohalide ions including thiocyanate (SCN^-^) and chloride (Cl^-^) to produce the oxidant acids hypothiocyanous acid (HOSCN) and hypochlorous acid (HOCl), respectively.^12^ Hydrogen peroxide is secreted by the airway epithelium and activated neutrophils, and both the Cl^-^ and SCN^-^ substrates are transported by CFTR into the airway surface layer.^13,14^ Though these highly potent oxidant acids are very effective in killing microbial pathogens, damage to the host tissues can also occur. MPO treatment has been shown to cause cell death of human airway epithelial (HAE) cells in submerged culture via oxidative stress.^2,15,16^ In addition, these studies have found that the addition of SCN^-^ and the subsequent generation of HOSCN reduces cytotoxicity dramatically in comparison to HOCl, which is highly cytotoxic. ^2,15^

SCN^-^ is drastically reduced in CF airways due to CFTR malfunction, potentially leading to worsened damage with increased generation of HOCl over HOSCN. ^16–18^ CFTR functional rescue by modulator drugs may increase the concentration of SCN^-^ in the airways, increasing the generation of HOSCN by MPO and leading to potential differences in mucociliary phenotypes. CFTR modulator medications, especially the highly effective triple combination therapy elexacaftor/tezacaftor/ivacaftor (ETI), provide ≥10% improvements in CFTR function *in vitro*.^17^ Though the effects of CFTR modulators on SCN^-^ concentration specifically in sputum samples has not been evaluated, a study that characterized SCN^-^ in the saliva of people with CF found that samples taken from people receiving single CFTR modulator therapies (such as ivacaftor, lumacaftor, or tezacaftor) had a significantly higher amount of SCN^-^ compared to those not taking modulators.^18^ It is likely with the introduction of the highly effective triple combination modulator therapy ETI that the amount of SCN^-^ in the airways is even further increased. While approximately 90% of the CF population is eligible for ETI, it remains inaccessible to a large portion of people with CF globally. A 2022 study estimated that only 12% of people with CF worldwide were receiving triple combination therapy.^19^ Furthermore, in the United States approximately ∼8% of people with CF are ineligible for CFTR modulators due to incompatibility with their genetic mutation and another ∼10% of people eligible for CFTR modulators are not using modulators due to various factors such as drug side effects.^20,21^ Given the disparities in access to modulator therapies, continued research into potential targets for anti-inflammatory treatments for those receiving and not receiving CFTR modulator therapies is merited.

Despite the potential clinical implications of increased MPO in causing oxidative stress and worsening of CF lung disease,^22^ it remains largely unknown how MPO affects MCT in the airways. Previous research found that MPO concentrations in people with non-CF bronchiectasis was correlated with more mucus plugging in patients, indicating presumably a lack of effective MCT.^23^ Prior *in vitro* studies evaluating the impacts of MPO on the airways have relied on cell lines that do not generate their own mucus and/or cilia, making evaluation of MCT impossible. Motivated by this, we used an *in vitro* model of the airway epithelium to study how MPO in the mucosal barrier affects mucociliary phenotypes. Specifically, we conducted these studies in BCi-NS1.1 human airway epithelial cultures fully differentiated at air-liquid interface which possess otherwise normal MCT function. We assessed these cultures after the exogenous addition of MPO in normal and CF-like conditions to determine their impact on MCT and airway surface liquid composition. Ultimately, these findings strengthen the rationale for therapeutically targeting MPO in CF and other muco-obstructive lung diseases.

## Results

Free MPO is released into the CF mucosal barrier via neutrophil degranulation and/or the degradation of NETs by DNase inhalation (**Figure 1A**). In designing our *in vitro* model to study how MPO affects mucociliary phenotypes, we created two different substrate solutions to react with MPO, each containing H_2_O_2_, SCN^-^, and Cl^-^ at concentrations relevant to the CF airway surface layer (**Figure 1B**).^24–27^ The only variable between the two substrate solutions was the concentration of SCN^-^. The substrate solution designated as “SCN^-^ high” contained the normal concentration of SCN^-^ ions found in the airway surface layer (500 µM) and the “SCN^-^ low” substrate solution contained 11 times less SCN^-^ (45.45 µM) to mimic the concentration in CF airways without CFTR modulators.^8,28,29^ The estimate of 45.45 µM was based upon a study using HAE cell cultures from CF donors grown at air-liquid interface (ALI) that found 11 times less SCN^-^ was transported from the media in the basolateral chamber into the apical airway surface layer compared to healthy controls.^29^ Based on the specificity constants, we calculated the theoretical ratio of HOSCN to HOCl production by MPO in the SCN^-^ high solution to be ∼27% HOSCN to ∼73% HOCl. In the SCN^-^ low solution, this ratio is calculated to be ∼80% HOSCN to ∼20% HOCl.^30^ We combined the substrate solutions with 30 µg/ml MPO, representative of the average concentration reported in CF sputum samples.^24,31^ To evaluate the specific contributions of MPO to mucociliary dysfunction, we also applied both the SCN^-^ high and low substrate solutions with no MPO added. We applied a 20 µL volume of the MPO-substrate or substrate-only solutions to cover the apical surface and allowed the cultures to incubate for 24 hours (**Figure 1B**). During this time, the small volume of liquid originally added to the airway surface layer was absorbed and mucociliary function could subsequently be evaluated.

**Figure 1.**
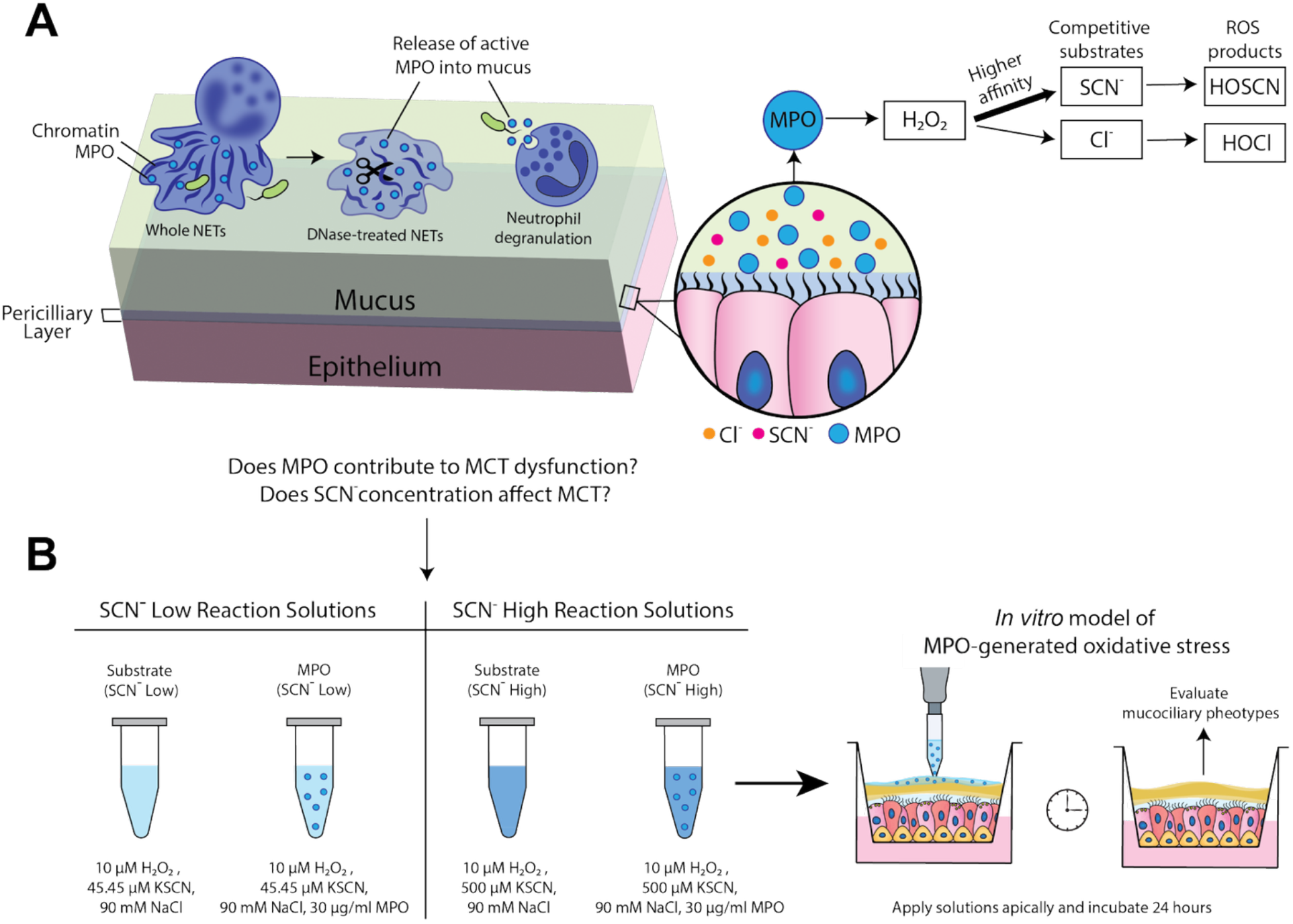
Modeling CF-like conditions *in vitro* to study the impact of MPO on mucociliary function. A) Schematic figure showing how neutrophilic inflammation in the airways leads to increased MPO concentrations in the mucus via degranulation and NETosis, and how MPO reacts with substrates in the mucus layer to form HOSCN and HOCl reactive oxygen species (ROS). B) Schematic figure showing the *in vitro* model created to address the central research questions of this work concerning the effects of MPO on MCT and the role of SCN^-^ concentration in altering mucociliary phenotypes. The SCN^-^ concentration was altered by creating two different substrate solutions, SCN^-^ low and SCN^-^ high. Four different treatment groups were established using the SCN^-^ low and SCN^-^ high substrate solutions, two of which contained MPO and two that consisted of the substrates alone. The substrate-only or MPO-substrate solutions were applied apically to the surface of HAE cultures and incubated for 24 hours.

### MPO reduces mucociliary transport *in vitro*

Particle imaging velocimetry was performed using fluorescence video microscopy of microspheres to determine the efficiency of MCT after treatment with either the MPO-substrate or substrate-only solutions for 24 hours (**Figure 2A**). MCT velocity was reduced most significantly in both MPO-substrate treatment groups regardless of SCN^-^ concentration (**Figure 2B, C)**. Both the substrate-only solutions alone had modest effects on the MCT velocity, though not significant from the untreated HAE cultures (**Figure 2B-C**). Based on the calculated velocities, the percentage of immobile microspheres was also determined, where a velocity of 0.1 µm/s was considered immobile (**Figure 2D**). This was chosen as the cut-off velocity because it meant that within the 20 seconds of the video, the microsphere failed to travel at least the length of 1 particle radius from its starting position. While an average of ∼40% of microspheres on the untreated control cultures were considered mobile, the vast majority (∼85-90%) of microspheres became immobile on the MPO-substrate treated cultures. To determine if the drop in MCT was due to changes to the ciliary beating, brightfield video was used to quantify the cilia beat frequency (CBF). Higher concentrations of hydrogen peroxide are known to reduce CBF transiently, however in this work the H_2_O_2_ concentration applied to the HAE cells was 10 µM, far below what is reported to significantly impair CBF or cause irreversible ciliostasis.^32,33^ No significant differences in the CBF between treatment groups 24 hours after treatment were seen (**Figure 2E**).

**Figure 2.**
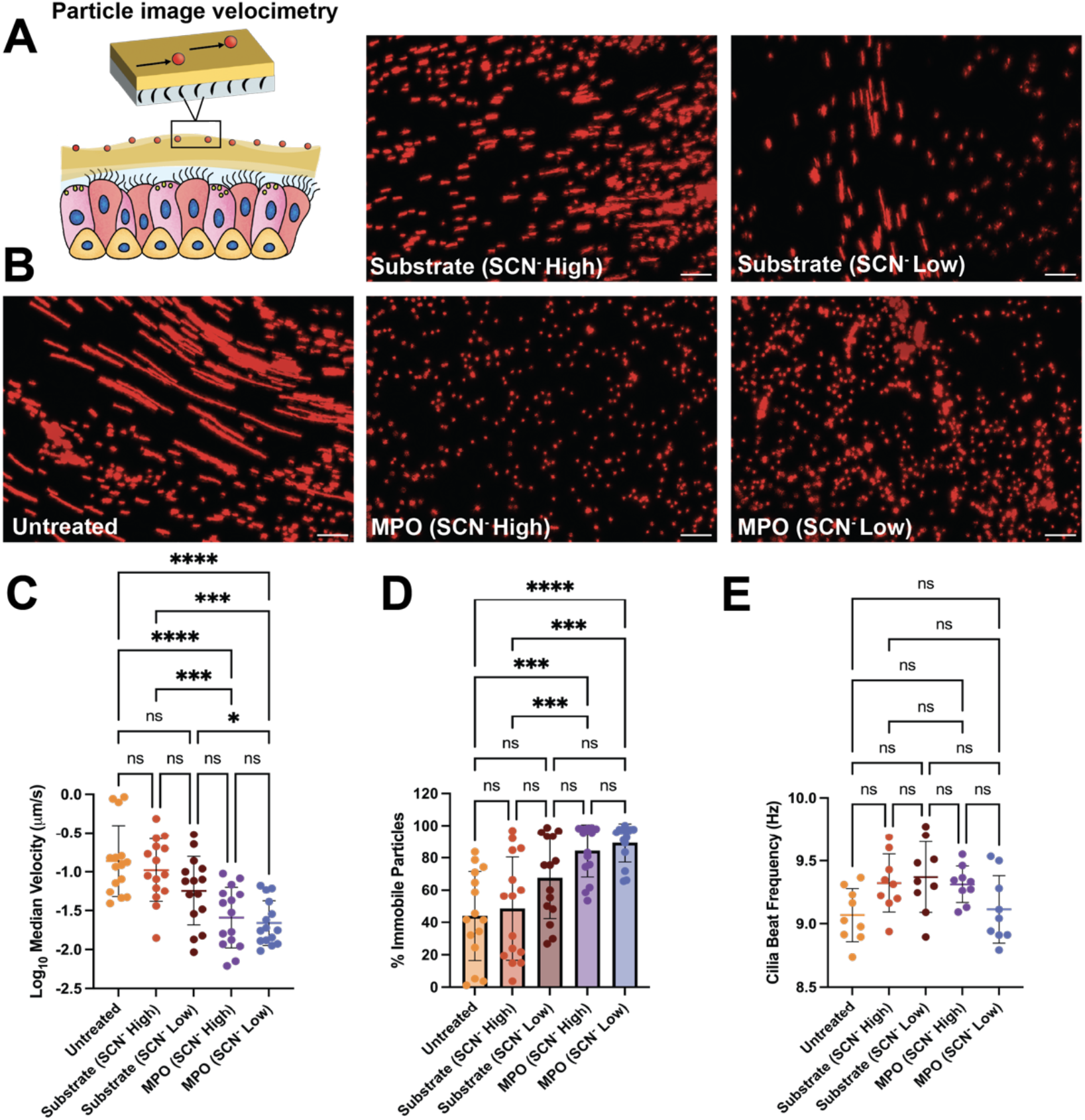
MPO reduces mucociliary transport *in vitro*. A) Schematic figure showing the data collection method for the evaluation of MCT using particle image velocimetry of fluorescent microspheres. B) Maximum pixel intensity projections created from videos of microspheres undergoing MCT across the apical surface of cultures after treatment with MPO-substrate or substrate-only solutions for 24 hours. Scale bars, 50 µm. C) The velocities of each individual microsphere were calculated using a MATLAB algorithm and log10 of the median velocity from each video was plotted. D) The percentage of microspheres considered immobile in each video (percent of microsphere trajectories with velocities under 0.1 µm/s). E) Cilia beat frequency of the HAE cultures was quantified using both FIJI and MATLAB. In panels C, D, and E the statistical analyses performed were one-way ANOVAs with Tukey’s multiple comparison tests (ns, no significance, * p < 0.05, *** p < 0.001, **** p < 0.0001).

We also investigated the viability and ciliation of the HAE cultures using flow cytometry. Loss of ciliation of the airway epithelium can occur as a result of exposure to ROS.^34^ Cells were stained with a viability dye and an antibody to detect alpha-tubulin, a cilia marker. We found no significant alterations in the percentage of dead cells in the total cell population (**Figure S1 A**). Though slightly decreased in all groups treated with ROS-containing solutions, the percentage of live ciliated cells in the total cell population did not exhibit major alterations (**Figure S1 B**) and was very similar to previous reports of percent ciliation of the BCI-NS1.1 cells.^35^ In addition, the mean fluorescence intensity of alpha-tubulin in the live alpha-tubulin positive cell population was also not changed significantly (**Figure S1 C**), indicating that there were no large changes to the expression of cilia on the identified ciliated population. We also measured transepithelial electrical resistance (TEER) and found an average of ∼10% decrease in TEER 24 hours after treatment of cultures with either of the MPO-substrate solutions and ∼5% average decrease in substrate-only treated cultures (**Figure S2**). This aligns with previous reports that MPO treatment can reduce the TEER in HAE cell monolayers. ^35,36^ The reduction in TEER seemed to indicate that the airway epithelium was still being affected by the MPO treatment despite the lack of changes to the cell viability, ciliation and ciliary function of the HAE cultures. Therefore, we investigated the composition of the airway surface layer was investigated for differences that could account for the large drop in MCT velocity.

### MPO alters the viscosity and macromolecular composition of airway surface liquid

The apical surfaces of the HAE cultures were washed with PBS following a 24-hour incubation with either the MPO-substrate or substrate-only solutions and the apical wash was collected (**Figure 3A**). We measured the viscosity of the apical wash using a microviscometer and found that the MPO-substrate treated apical washings displayed the highest viscosity, suggesting that the macromolecular content was increased in these samples compared to the untreated samples (**Figure 3B**). The MPO/SCN^-^ low treatment group had the highest viscosity. In addition, we determined that exogenous treatment of mucus harvested from HAE cultures with MPO did not significantly alter its viscosity (**Figure S3**). To detect changes in the macromolecules produced in response to MPO exposure, the total protein and DNA content of the apical surface washes were quantified. The total protein in the whole apical wash of all the MPO-substrate and substrate-only treated cultures was significantly higher than the untreated cultures (**Figure 3C**). The protein concentration was highest in the MPO-treated SCN^-^ high HAE cultures (**Figure 3C**). In quantifying the percent change in total protein compared to the untreated cultures, we found that there were significant increases for the SCN-low substrate and MPO SCN^-^ low conditions (**Figure 3D**). The DNA was found to be highest in the apical wash from the MPO/SCN-low substrate treatment, with it being the only treatment to show statistical significance in the percent change over the untreated condition (**Figure 3E**). We confirmed this using another DNA staining method by adding Sytox Green to the apical wash samples (**Figure S4**). The increase in DNA could indicate cytotoxicity in response to the upregulated production of HOCl. However, flow cytometry analysis of the HAE cells revealed no significant changes in the viability **Figure S1A**. We also evaluated cellular viability based on lactose dehydrogenase (LDH) activity in the basolateral media from each individual culture 24 hours pre-treatment and 1 hour after treatment and calculated the percent change (**Figure S5**). The MPO/SCN^-^ low treatment group was the only one that exhibited increases in the percent change of LDH activity above the untreated control. Both the increased DNA content in the airway surface layer and the minor increase in LDH activity suggest very small and difficult-to-detect enhancements in cell death.

**Figure 3.**
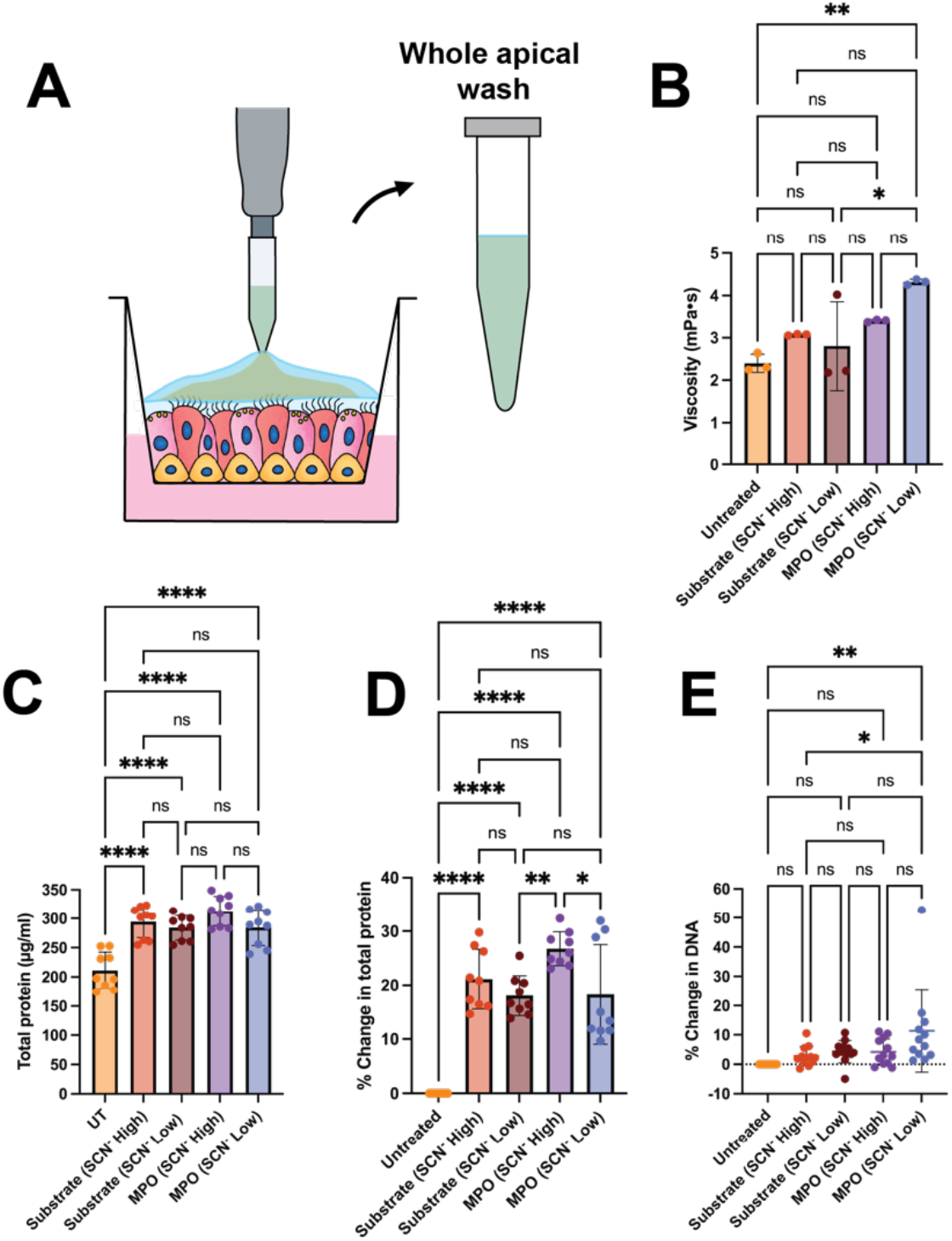
MPO alters the viscosity and macromolecular composition of airway surface liquid. A) Schematic figure showing the collection of the whole apical wash from HAE cultures. B) Viscosity of the apical wash samples from each treatment group quantified using a microviscometer. C) The total protein in the apical wash was quantified using a bicinchonic acid based colorimetric reaction. The concentration of total protein in each sample was determined using a standard curve of bovine serum albumin and D) the percent change in the absorbance values was calculated in relation to the untreated group. E) The DNA in the apical wash quantified using 3,5-diaminbenzoic acid (DABA) to produce a fluorescent reaction. The percent change in the fluorescence was calculated compared to the untreated group. Statistical significance in B, C, D and E was determined using one-way ANOVAs with Tukey’s multiple comparison tests (ns, no significance, * p < 0.05, ** p < 0.01, **** p < 0.0001).

### MPO does not alter mucin expression or mucin-mucin crosslinking

Based on the finding of upregulated protein expression in the apical wash, we investigated if the treatments caused alterations to mucin glycoprotein expression and network structure of mucus. It should be noted as well a prior study characterized the effects of applying exogenous hydrogen peroxide to HAE cells on the expression of multiple mucin types in the airways and found that only MUC5AC mRNA expression was increased. However, the range of hydrogen peroxide concentrations tested were 10-100 times higher than this study.^36^ MUC5AC is generally known to form more viscoelastic mucus gels when overexpressed that inhibit mucociliary transport significantly.^37^ Though MUC5AC is known to be the mucin typically upregulated by ROS and other neutrophil inflammatory mediators, MUC5B may also be upregulated. MUC5B, the other major secreted mucin in the airway, is known to be overexpressed in CF often due to the cycle of “sterile inflammation.” Highly viscoelastic mucus causes necrosis of airway epithelial cells and leads to increased secretion of IL-1β that induces secretion of MUC5B.^38^ Considering this, we assessed the total mucin content of the airway surface layer after MPO-substrate or substrate-only treatment. To detect mucin glycoproteins in the airway surface layer, we took the whole apical wash from HAE cultures and isolated the >100 kDa fraction using centrifugal filtration (**Figure 4A**). Mucins are very large proteins (2 to 50 MDa) that will be retained in the >100 kDa fraction, while smaller glycoproteins will be removed.^3^ We then took the >100 kDa fraction and fluorometrically detected O-linked glycosylation to evaluate mucin content (**Figure 4B-D**). We found no significant differences, indicating no changes in mucin expression at the protein level. To ensure there were no changes at the mRNA level, we also evaluated the expression of MUC5AC mRNA using quantitative PCR (qPCR). In agreement with the mucin detection assay, there were no significant changes found in MUC5AC mRNA expression (**Figure S5**). In addition to mucin expression, we wanted to investigate changes to the microstructure of mucus. The microstructure of the mucus gel is directly influenced by disulfide bonding occurring between thiol groups in cysteine-rich regions of the mucin proteins. We quantified the disulfide bonds within the >100 kDa, mucin containing fraction of the apical wash in HAE cells treated with the substrate or MPO-substrate solutions using a fluorometric assay (**Figure 4E-G**). We found no significant differences between the disulfide bonds in groups, suggesting that MPO does not catalyze formation of additional mucin-mucin crosslinks at the concentrations tested in this study.

**Figure 4.**
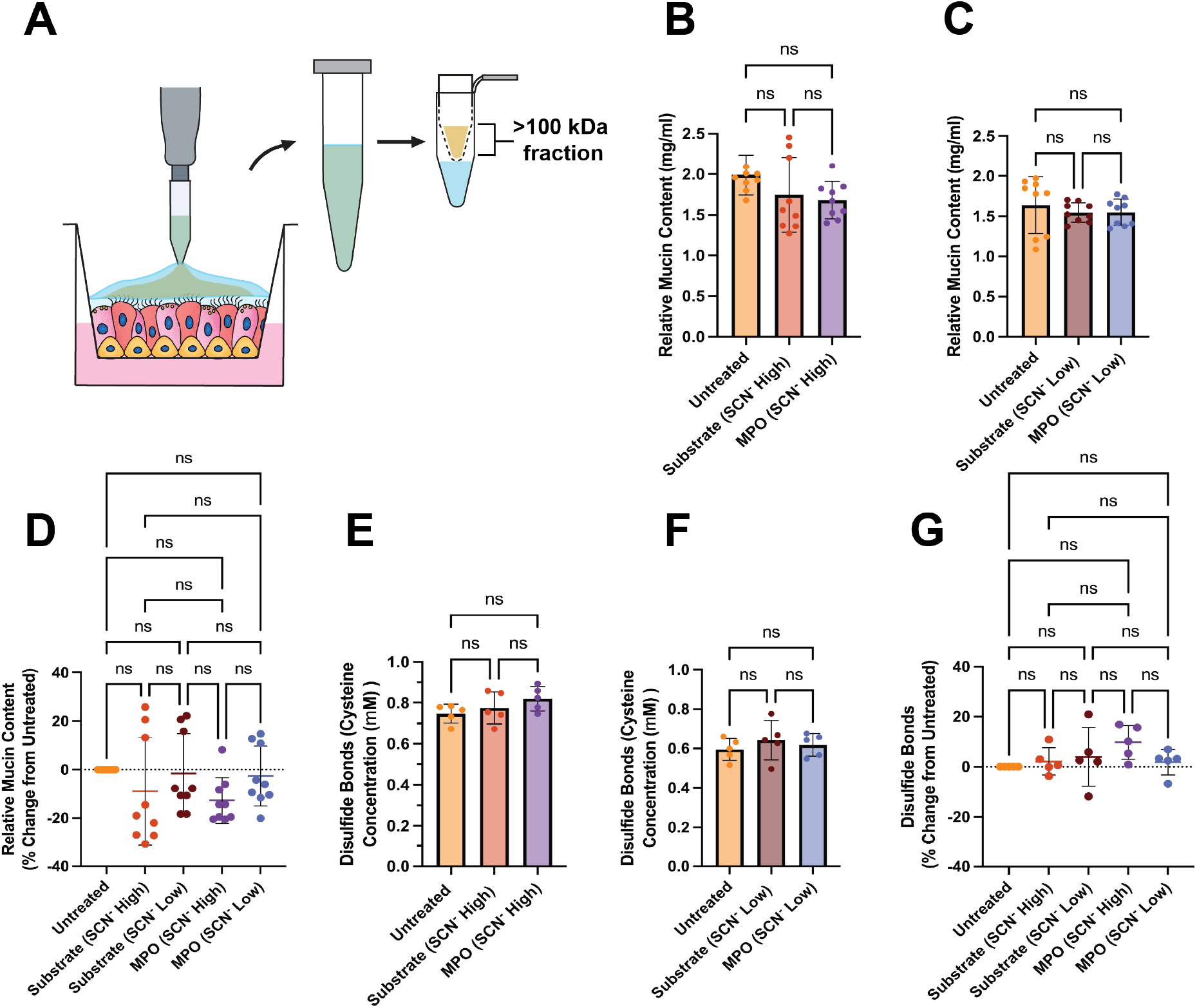
MPO does not alter mucin expression or mucin-mucin crosslinking. A) Schematic figure showing the collection of the whole apical wash from HAE cultures and subsequent separation of the >100 kDa fraction using a centrifugal filter. B, C) The relative mucin content of the >100 kDa fraction was assessed using a 2-cyanoacetamide based reaction with O-linked glycans on mucins. A standard curve of bovine submaxillary gland mucins was used to calculate the relative concentration of mucins in the samples and D) the percent change in fluorescence values was calculated in comparison to the untreated group. E, F) The disulfide bonds in the >100 kDa fraction were quantified using by blocking free cysteines, followed by treatment with a reducing agent to expose only the disulfide-bonded cysteines. Cysteines were detected fluorescently with monobromobimane. The concentration of disulfide bonds in samples was determined using a standard curve of L-cysteine and G) the percent change in the fluorescence values was calculated in relation to the untreated group. Statistical significance in panels B-G was determined using a one-way ANOVA with Tukey’s multiple comparison tests (ns, no significance).

### MPO impairs MCT similarly to NE

We next compared the effects of MPO to neutrophil elastase (NE), another neutrophilic granular protein found in abnormally high concentrations in CF sputum. NE is a serine protease that has been widely studied and is well known to induce MUC5AC secretion by HAE cells.^39–41^ NE has been associated with worsening of lung function and airway clearance in CF. ^31,42–44^ A previous study characterizing the effects of NE on MCT was done on anesthetized quails, in which there was a reduction in MCT over the course of 60 minutes after treatment with 100 µg/ml and 300 µg/ml NE.^45^ This was largely attributed to increased protein and DNA macromolecules found in the tracheal lavage fluid. In addition, aerosolized NE was shown to reduce MCT in ovine airways after 6 hours.^46^ However, the effects of NE on MCT on human airway tissue cultures *in vitro* have not been directly characterized to the best of our knowledge. To compare the impact of MPO and NE on MCT, we added either 30 µg/ml MPO or 25 µg/ml NE, representative of the average concentrations of each found in CF sputum samples, to the SCN^-^ low substrate solution.^24,31,47^ By comparison, healthy controls were found to have ≤0.1 µg/ml of MPO and NE in sputum samples.^24,31,47^ We found that both NE and MPO significantly reduced MCT velocity compared to the untreated and SCN^-^ low substrate solution-treated groups alone (**Figure 5A, B**). MPO exhibited the lowest median MCT velocities that were significant from NE, though the percentage of immobile microspheres was not significant between the NE and MPO treated cultures (**Figure 5B, C**). We also evaluated the CBF and found similar results to the MCT experiments done in Figure 2E. The substrate treated cultures had a slightly increased CBF, while the NE treated cultures had a lower CBF. This is consistent with some literature implying that NE may slow ciliary beating, especially at higher concentrations.^48^

**Figure 5.**
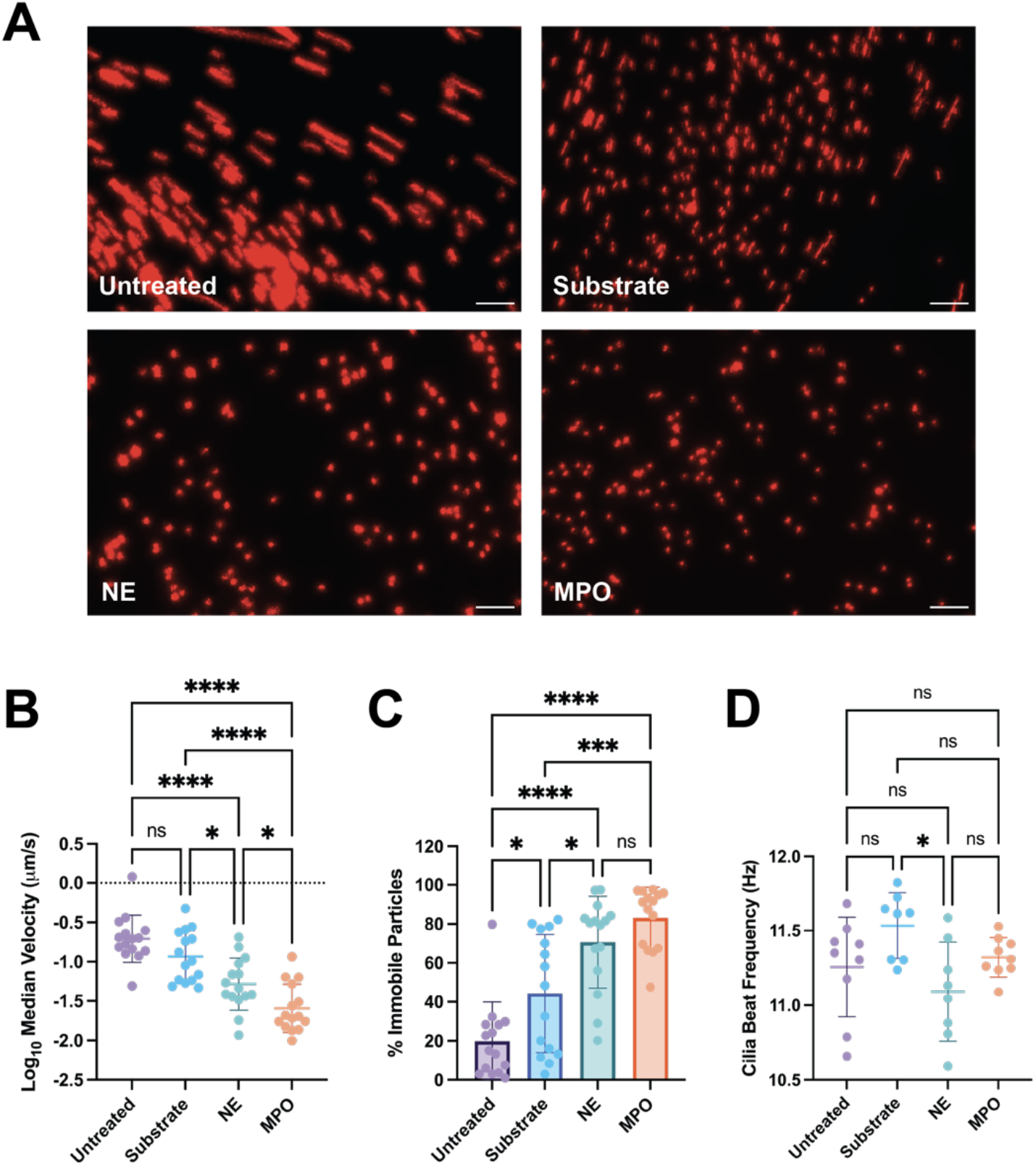
MPO impairs MCT similarly to NE. A) Maximum pixel intensity projections of microspheres being transported over the apical surface of HAE cultures after treatment with MPO-substrate, substrate-only, or NE-substrate solutions for 24 hours. Scale bars, 50 µM. B) The MCT videos were analyzed with a MATLAB algorithm to determine the velocities of each microsphere and the log10 of the median velocity for each video was plotted. C) Microspheres were considered immobile if their velocity was less than 0.1 µm/s. The percentage of immobile particles was quantified and plotted for each video taken. D) The cilia beat frequency of the HAE cultures was quantified using FIJI and MATLAB analyses. One-way ANOVAs with Tukey’s multiple comparison tests were used to determine statistical significance in B, C, and D (ns, no significance, * p < 0.05, *** p < 0.001, **** p < 0.0001).

## Discussion

We found that the application of exogenous MPO, likely to be at elevated levels due to chronic inflammation in the CF lung, significantly slowed MCT in differentiated HAE cultures. It should be noted excessive inflammation in the airways, primarily driven by neutrophils, remains a problem for many people with CF with or without access to modulator therapies. Recent studies have found that CFTR modulators are able to only partially dampen inflammation. One study found that inflammatory markers in CF sputum samples were reduced after ETI therapy to be very similar to people with non-CF bronchiectasis.^31^ Another recent study found that inflammatory markers in the sputum of CF patients taking ETI were decreased to levels similar to what is found in samples from people with primary ciliary dyskinesia.^49^ In both cases the sputum inflammatory markers in people with CF taking CFTR modulator drugs are decreased to be comparable with other chronic lung diseases, yet still much higher than healthy controls. Considering this, MPO-mediated defects in airway clearance due to neutrophilic inflammation could reduce lung function in individuals of CF independent of their use of highly effective CFTR modulators.

Within the airway surface layer in the CF-affected lung, the high extracellular concentrations and elevated enzymatic activity of MPO is due to increased neutrophil degranulation and NETosis. CF neutrophils have been shown to have an increased propensity to undergo degranulation compared to healthy neutrophils, releasing free MPO into the airway surface layer.^50^ In addition, very high amounts of NETs are released into the mucus layer in CF.^51^ Multiple studies have shown that MPO remains enzymatically inactive in NETs due to being complexed with DNA.^10,11^ However, daily inhalation of DNase is one of the most widely used CF treatments to degrade the DNA ultrastructure of NETs and improve MCT efficiency.^21,52^ Previous research has found that DNase treatment of NETs causes the release of active MPO.^11^ Therefore, our results showing that MPO decreases MCT supports the notion that there may be unintended effects of DNase treatment in releasing active NET-bound MPO.

Both Cl^-^ and SCN^-^ are transported via CFTR and the balance of these ions cause MPO to generate different oxidant products. Without CFTR modulators, SCN^-^ concentrations in CF airways are very low, likely encouraging the production of the highly damaging oxidant HOCl.^8^ It is plausible that people taking CFTR modulator therapies may have concentrations of SCN^-^ restored to near normal levels and enhanced HOSCN production in the airways.^18^ However, we found MCT is significantly impaired by MPO treatment independent of SCN-concentration. After the 24-hour treatment with MPO, we found no major changes in ciliation or cilia beat frequency, suggesting the inhibition of MCT was caused by changes to the airway surface layer. After washing the apical surface layer off each of the cultures and measuring the viscosity of each, we found that they were highest in the MPO-treated cultures. Specifically, the MPO-SCN^-^ low substrate solution condition had the highest viscosity. Therefore, we explored the macromolecular composition of the apical surface layer of each treatment group. We found that MPO in combination with both SCN^-^ high substrate solution resulted in the highest protein content in the airway surface layer. It is unlikely the apparent increase in concentration is due to the exogenous addition of MPO alone as we estimate this would yield <1% increase in protein given the total effective dose of MPO being 0.6 µg per culture. Therefore, increases in protein content of airway surface liquid following MPO-substrate and substrate-only treatments are presumably due to elevated secreted protein concentrations.

In addition to changes in protein content of the mucus, treatment with MPO and SCN^-^ low substrate in combination lead to the highest increases in DNA content in the apical surface. This could be because of the increased production of HOCl, which is known to be more cytotoxic over HOSCN. However, the mucus layer does seem to be protective against oxidative stress, as we did not find that our MPO-substrate treatments were highly cytotoxic. This is in agreement with prior studies that have shown that the components of the extracellular matrix (ECM) and CF mucus interact strongly with MPO likely due to the interaction of cationic MPO with anionic macromolecules such as DNA, glycosaminoglycans, and mucins.^10,16,53,54^ Multiple studies have concluded that MPO often reacts primarily with the components of CF mucus rather than the underlying cells. In one study, Chinese hamster ovary cells were treated with 66 µM MPO and 300 µM H_2_O_2_, which caused ∼80% of cells to die. When heat inactivated CF sputum was added to the media at a 1:20 dilution and treated again with MPO and H_2_O_2_, the cytotoxicity dropped to ∼0%.^10^ This is supported by other research that conducted analysis of proteins in both CF sputum and bronchoalveolar lavage samples and found hallmarks of HOCl-mediated oxidation.^8,16^ The antioxidant molecule glutathione (GSH) is also secreted by the airway epithelium to scavenge oxidants in the mucus. A previous study using 16HBE14o-cells mounted in an Ussing chamber found that the addition of physiologically relevant concentrations of glutathione completely prevented HOCl-induced disturbances in the transepithelial resistance.^55^ Therefore, it is plausible that we could not detect major differences in viability. Though we did not see large changes in the cytotoxicity of our MPO-substrate treatments, we still believe that the elevated DNA found in the apical surface layer is from very minute increases in cell death that are not easily detectable. DNA is a large macromolecule and even slight increases can contribute largely to enhanced mucus viscoelasticity.^56^

Given the increase in protein content of the apical wash and the ability of many inflammatory mediators to induce mucin protein secretion in CF, we also evaluated mucin expression after MPO treatment. However, we found no upregulation of mRNA or protein expression for mucins. In addition, we assessed the formation of disulfide bonds between mucins after MPO treatment given that HOCl can oxidize thiol groups. We did not find any changes in disulfide bond formation between mucins, indicating that the increased viscosity of mucus was not due to increased mucin crosslinking. Similar peroxidase enzymes secreted by the airway epithelium, thyroid peroxidase (TPO) and lactoperoxidase (LPO), were shown to enhance disulfide bonding between thiolated polymers.^57^ In these experiments, the mucus layer of HAE cultures was completely removed using multiple dithiothreitol and PBS washes, and a solution of the thiolated polymer was overlaid on the apical surface of the HAE cultures. Conversely, we find no evidence that MPO treatment alters mucin expression or crosslinking. It should be noted HAE cells express other proteins with antioxidant activity such as glutathione peroxidase, catalase, glutathione S-transferase, peroxiredoxin, or thioredoxin which may have had a protective effect and limited MPO-mediated changes to mucin content or cross-linking. ^14,55,56^ As such, we would speculate MPO-mediated deficits in MCT could, at least in part, be a result of disrupted epithelial barrier integrity, as evidenced by changes in TEER, which may cause fluid and ion imbalance leading to dehydration of the airway surface liquid layer. This could in effect increase the concentration of mucin and other apically secreted proteins without having a direct impact on the extent of mucin glycoprotein expression.

We also compared the effects of MPO on MCT to another neutrophilic granular protein that is very well characterized to induce mucin secretion and deficits in MCC. NE is a potent inducer of MUC5AC expression by the airway epithelium. We applied MPO and NE at concentrations similar to their reported average concentrations in CF sputum samples, 30 µg/ml and 25 µg/ml respectively.^24,31,47^ We found that MCT was significantly impaired by both NE and MPO, with MPO having the lowest MCT velocities compared to NE. One contributing factor to this result may be that NE is a protease that cleaves various proteins in the airway surface layer, potentially loosening the mucus gel network and enhancing MCT. Overall, our data indicate both MPO and NE are important NET-associated components to consider in CF and other chronic lung conditions where mucociliary clearance is impaired.

This work establishes MPO can directly impair MCT *in vitro* via cell-mediated changes to airway surface liquid properties. Adjusting the SCN^-^ concentrations to generate primarily HOSCN over HOCl did not make substantial differences in MCT. MPO showed a similar decrease in MCT to NE, another granular protein found to be upregulated in CF airways and known to cause deficits in airway clearance function. We also acknowledge the limitations of this study, including the use of BCI NS1.1 immortalized normal HAE cell line instead of primary CF cultures. In addition, we did not consider the use of hypertonic saline in our experiments, which is inhaled daily by many people with CF and can raise the amount of chloride in the airways. This may boost HOCl production even further and contribute to more damage. These data show MPO as a potential therapeutic target to improve MCT and further reduce mucus plugging for people with CF both receiving and not receiving CFTR modulator therapies. The MPO-mediated reduction in MCT could also apply to similar lung pathologies with neutrophil-driven inflammation such as non-CF bronchiectasis, chronic obstructive pulmonary disease (COPD), or acute infections.

## Methods

### Air Liquid Interface Culture of Human Airway Epithelial Cells

The h-TERT immortalized human airway epithelial basal cell line, BCI-NS1.1 (provided by Ronald Crystal, Cornell), was cultured in a flask in Pneumacult Ex-Plus media (Stemcell Technologies) until cells reached 70-80% confluency. To establish air-liquid interface (ALI) cultures, cells were seeded at a density of 10^4^ cells/cm^2^ on collagen-coated 6.5 mm Transwell cell culture inserts (Corning) with Pneumacult Ex-Plus media added to both the apical and basolateral compartments. After the cells formed a confluent monolayer, the media was removed from the apical compartment to expose cells to the air. The media in the basolateral compartment was also switched to Pneumacult ALI (Stemcell Technologies) once airlifted. The cells were cultured at ALI for at least 28 days to ensure full differentiation of the epithelium, and media was changed every two days. After reaching maturity, mucus was washed from the surface of the cells at least once per week by adding 250 µL of DPBS (without Ca^+^ and Mg^2+^) to the apical compartment. The cells were incubated with the PBS for 30 minutes at 37°C and 5% CO_2_, then the PBS was removed by aspirating or by collecting with a pipette if using the mucus washings.

### MPO/NE/Substrate Solution Treatment of HAE Cultures

All HAE cells were washed 24 hours prior to treatment with MPO, NE or substrate solutions. Potassium thiocyanate (KSCN) and sodium chloride (NaCl) were dissolved in sterile ultrapure water. Hydrogen peroxide was diluted in sterile ultrapure water. The NaCl, KSCN, and H_2_O_2_ solutions were combined into one solution and sterile filtered. MPO from human neutrophils (Athens Research and Technology) or a vehicle (substrate only) was added to each solution to achieve a final concentration of 500 µM KSCN, 90 mM NaCl, and 10 µM H_2_O_2_ in the SCN^-^ high substrate solution and 45.45 µM KSCN, 90 mM NaCl, 10 µM H_2_O_2_ in the SCN^-^ low substrate solution. In solutions containing MPO, the final concentration of MPO was 30 µg/ml. In solutions containing NE from human neutrophils (Athens Research and Technology), the final concentration of NE was 25 µg/ml. Once the solutions were prepared, fully differentiated HAE cells were treated apically with 20 µL of each mixture for 24 hours. During the 24-hour incubation period, the 20 µL of liquid applied to the surface of the cultures was completely absorbed.

### Mucociliary Transport Quantification

All HAE cultures were washed and treated with MPO, NE or substrate solutions as described above. MCT was quantified using particle image velocimetry data collection and analysis methods described previously.^58^ 2 µm red fluorescent microspheres (Thermo Fisher FluoSpheres) were sonicated for 10 minutes, then diluted 1:3000 in sterile PBS and mixed thoroughly. 20 µL of the microsphere solution was applied apically to the mucosal layer of the cells. Cultures incubated with the microspheres for 15 minutes at 37°C, 5% CO_2_. After this incubation, the cultures were removed and placed in a 6 well plate using tweezers so the cell culture insert membrane was flush with the bottom of the 6 well plate and could be imaged clearly. This process of applying the microspheres, incubating for 15 minutes, and moving the cultures to was repeated for each batch of 3 cultures in each treatment group immediately prior to imaging. The apical surface of the cultures were imaged using fluorescence video microscopy. Videos of microspheres undergoing MCT in 5 random regions of each HAE culture were taken at 10x magnification for 20 seconds at an exposure time of 150 ms. Each video was analyzed to quantify the velocity of the microspheres using MATLAB. The MATLAB analysis identified and tracked the trajectories of the microspheres, then computed the velocity of each based on the x and y coordinates of the microspheres’ trajectory. The median MCT velocity from each video was plotted.

### Cilia Beat Frequency Quantification

The cilia beat frequency was quantified using analysis of brightfield microscopy videos using previously disseminated methods.^58^ Directly after taking the MCT videos, the microscope settings were switched to brightfield. The focus on the epithelium was adjusted until ciliary beating was clearly visible. Videos were taken in 3 random areas of each HAE culture using 10x objective for 10 seconds at 20 ms. The CBF was quantified using FIJI and MATLAB to analyze each video. Briefly, the mean intensity over the time of the video was obtained in FIJI in 3 different randomly selected regions of each video. The mean intensity versus time data was saved and analyzed using a MATLAB script. The MATLAB script identified the local maxima that represent the beating of cilia in the mean intensity versus time data. The number of local maxima over the time was computed and yielded the cilia beat frequency of each of the 3 regions selected from each individual video. The mean CBF value from each video was plotted.

### Protein quantification in HAE Apical Wash

The total protein in the apical wash of HAE cells was determined using a bicinchonic acid (BCA) assay (Thermo Fisher Scientific), according to kit instructions. HAE cells were washed according to the methods described above following 24 hours of treatment with MPO-substrate or substrate-only solutions. The whole apical wash collected from 2 cultures in each treatment group were pooled and thoroughly mixed. A standard curve was prepared using dilutions of bovine serum albumin, and 25µL of samples and standards were added to a clear 96 well plate. 200 µL of the protein detection reagent was added to the wells and incubated at 37°C for 30 minutes to allow the colorimetric reaction to take place. The absorbance was read on a plate reader at a wavelength of 562 nm.

### DNA Quantification in HAE Apical Wash

The total DNA in the apical wash of the HAE cells was determined using 3,5-diaminbenzoic acid (DABA), which produces fluorescent compound when reacting with the aldehydes in DNA.^59^ The whole HAE apical washes from 2 cultures in each treatment group were pooled and thoroughly mixed. 30 µL of the apical wash samples were mixed with 30 µL of 20% w/v DABA and incubated for 1 hour at 60°C on a heat block. The reaction was quenched with 1 mL of 1 M HCl and mixed. Samples were added to the wells of a black 96 well plates and the fluorescence was read at excitation/emission wavelengths of 340/420 nm.^59^

### Viscosity Quantification of Apical Wash

After 24 hours of treatment, the whole apical wash from 2 cultures in each treatment group were collected, pooled, and mixed thoroughly. The viscosity of the apical wash samples from each condition were quantified at room temperature using the Rheosense microviscometer with an A05 chip installed. A viscosity standard of 10 mPa•s was tested prior to testing samples to ensure that the microviscometer was properly functioning. Prior to testing samples, the microviscometer was run on cleaning mode twice with deionized water. The microviscometer pipette was loaded with ∼400 µL of the apical wash samples, ensuring that any air bubbles were removed. The viscosity was measured using the automatic mode, which determines the optimal parameters for microviscometer sample priming and measurements including the volume and shear/flow rate.

### Mucin Content Assay

The whole apical wash was collected from 2 HAE cultures in each treatment group and mixed thoroughly. 450 µL of the whole apical wash was added to a 0.5 mL 100 kDa molecular weight cut-off Amicon Ultra centrifugal filter (Sigma Aldrich) and centrifuged for 20 minutes at 14,000xg. The centrifugal filter was flipped upside down into a new microcentrifuge tube and the contents was removed by centrifuging for 2 minutes at 2000xg to yield the >100 kDa fraction of the HAE apical wash. The mucin content of the samples was evaluated using a previously established fluorometric assay to detect the O-linked glycans of glycoproteins.^60^ Briefly, a standard curve of bovine submaxillary gland mucins (Sigma Aldrich) was made using serial dilutions of a 2 mg/ml stock in PBS. a solution of 0.6 M 2-cyanoacetamide (CNA) was mixed with a solution of 0.15 M sodium hydroxide (NaOH) to achieve final concentrations of 0.1 M CNA and 0.025 M NaOH. 60 µL of this reagent was added to 30 µL of each sample and incubated on a heat block at 100°C for 30 minutes. The tubes were centrifuged briefly using a tabletop centrifuge before quenching the reaction. The reaction was quenched by adding 500 µL of a buffer consisting of 0.6 M boric acid with a pH of 8.0 to each sample. The samples were added to a black 96 well plate and the fluorescence was quantified using a plate reader with excitation/emission wavelengths of 336 and 383 nm, respectively.

### Disulfide bond quantification

The whole apical wash was collected from 2 HAE cultures from each treatment group after 24 hours and mixed thoroughly. 450 µL of the whole apical wash was added to a 0.5 mL 100 kDa molecular weight cut-off Amicon ultracentrifuge filter (Sigma Aldrich) and centrifuged for 20 minutes at 14,000xg. The disulfide bonds were quantified using a previously established fluorometric assay.^24^ Briefly, 50 µL of samples were aliquoted and 8 M Guanidine-HCl was added to achieve a final volume of 500 µL. The diluted samples were treated with 10% v/v 500 mM iodoacetamide (final concentration of 50 mM) for 1 hour at room temperature to block free cysteines. Samples were then treated with 10% v/v 1 M dithiothreitol (DTT) at 37°C for 2 hours to reduce disulfide bonds. Using 7 kDa molecular weight cut off Zeba desalting columns according to manufacturer’s instructions (Thermo Fisher Scientific), small molecules were removed from the samples and buffer exchange was performed with 50 mM Tris-HCl pH 8.0. A standard curve of L-cysteine was made using serial dilution of a 2.5 mM stock in Tris-HCl buffer. 70 µL of each sample or standard was loaded into black 96 well plates, then an equal volume of 2 mM monobromobimane (Cayman Chemical) was added to each well to fluorescently detect cysteine residues. The plate was incubated for 15 minutes covered at room temperature, then the fluorescence was quantified using a plate reader with an excitation/emission wavelength of 395 and 490 nm, respectively.

## Acknowledgments

This project was funded by the Cystic Fibrosis Foundation (DUNCAN24G0), the National Institutes of Health (R01 HL160540, F31HL176146).

