## Supplementary Information for "Myeloperoxidase impairs mucociliary transport on human airway epithelium"

### Supplementary Methods

#### Flow Cytometry of HAE Cells

Cells were washed with PBS twice to remove the mucus. Trypsin was added to the apical and basolateral chambers and incubated until cells were rounded and dissociated from the transwells. The cells from each individual culture were collected in separate tubes containing a trypsin neutralizing solution. The cell suspension was centrifuged and resuspended in PBS, then strained twice using a 40 μM cell strainer to break up clumps of cells. The cells were washed in PBS, then resuspended in ice cold FACS buffer (2% heat inactivated FBS in PBS) containing Trustain FcX (Biolegend) diluted 1:100 to prevent non-specific antibody binding. After blocking for 15 minutes at 4°C, cells were washed in ice cold PBS. Cells in the dead control group were heat killed by placing the tube on a heat block at 100°C for 5 minutes. Samples were stained with a solution containing a combination of Zombie Fixable NIR dye (Biolegend) diluted 1:1000 to detect viability and an AlexaFluor 488 conjugated anti-alpha tubulin diluted 1:40 to distinguish ciliated cells (BioLegend #627906). Single stain controls and unstained samples were also included for compensation and gating. After staining for 30 minutes covered on ice, cells were pelleted and resuspended in 4% paraformaldehyde. Cells were incubated in the paraformaldehyde fixative for 20 minutes at 4°C, then washed and resuspended in FACS buffer. Cells were run on the flow cytometer (BD Biosciences FACS Celesta). To gate populations, doublets and debris were excluded. The singlet cell population was gated based on unstained and single stained controls to examine cell viability and percentage of live ciliated cells in the total HAE cell population. The mean fluorescence intensity of the live alpha-tubulin positive singlet cell population was also plotted to evaluate expression of cilia.

#### Transepithelial Electrical Resistance (TEER) Measurement

The pre-treatment TEER of cells was taken directly before treatment with MPO-substrate or substrate-only solutions. To prepare fully differentiated HAE cells for TEER measurements, the basolateral media was removed and replaced with 1 mL of room temperature PBS. 500 μL of room temperature PBS was also added to the apical compartment, then cells were incubated for 15 minutes at room temperature. During this incubation, the TEER probe was sterilized in 70% ethanol. After 15 minutes, the probe was rinsed in room temperature PBS. To conduct the measurements, the probe was held at a 90° angle and three measurements were taken around three different regions of each culture. After 24 hours of treatment with either MPO-substrate or substrate-only solutions, the measurements were repeated, with four cultures in each treatment group.

#### Multiple Particle Tracking Microrheology

For microrheology experiments, 20 μl of sample was added to a microscopy chamber created by sticking a vacuum-grease covered O-ring to a glass microscope slide. 1 μl of PEG-coated PS nanoparticles diluted to 0.0025% w/v in ultrapure water was added to the sample, then the microscopy chamber was sealed using a cover slip. Samples were incubated at 37°C for 2 hours, then imaged using fluorescence video microscopy. Nanoparticle diffusion was imaged using 63x magnification with a water-immersion objective on a Zeiss LSM 800 microscope. 10 second videos of the fluorescent nanoparticles were captured within each hydrogel at a frame rate of 33 Hz in 5 different regions of the sample. Using the videos of nanoparticle diffusion, the MSD as a function of the lag time (τ) was quantified for each particle in MATLAB using the equation $\langle \Delta r^2(\tau) \rangle = \langle (x^2+y^2) \rangle$.

**Quantification of DNA in Apical Wash via Sytox Green Fluorescence**
Apical wash samples were collected and thoroughly mixed, then added to a black 96 well half area plate. Sytox green (Thermo Fisher Scientific) DNA stain was diluted 1:5000 in PBS, then 10% v/v was added to the apical wash samples in each well. The plate was incubated covered for 15 minutes, then was read using a plate reader at excitation/emission wavelengths of 504/523 nm to quantify Sytox green fluorescence. The percent change in relative fluorescence units of each group was calculated compared to the untreated sample.

**Cytotoxicity**
50 μL of the basolateral media was sampled from each culture 24 hours prior to treatment to establish a baseline level of LDH for each culture. LDH activity was measured using an assay kit according to manufacturer's instructions (Promega). An equal volume of reaction solution was added to the media samples in a clear 96 well plate and incubated covered at room temperature for 30 minutes. 50 μL of stop solution was added to quench the reaction and the absorbance was read at 490 nm on a plate reader. The LDH activity assay was repeated 1 hour after treatment with MPO-substrate or substrate-only solutions and the percent change from the baseline LDH levels for each culture were calculated.

**MUC5AC Quantitative PCR**
Following treatment with either the substrate only or MPO-containing solutions for 24 hours, RNA was extracted from the cells using the RNeasy kit (Qiagen) according to manufacturer's instructions. cDNA was generated from the extracted RNA using SuperScript III (Invitrogen). Quantitative PCR was performed using PowerUP SYBR Green Master Mix (Applied Biosystems) and forward and reverse primers specific for MUC5AC (Integrated DNA Technologies).

### Supplementary Figures

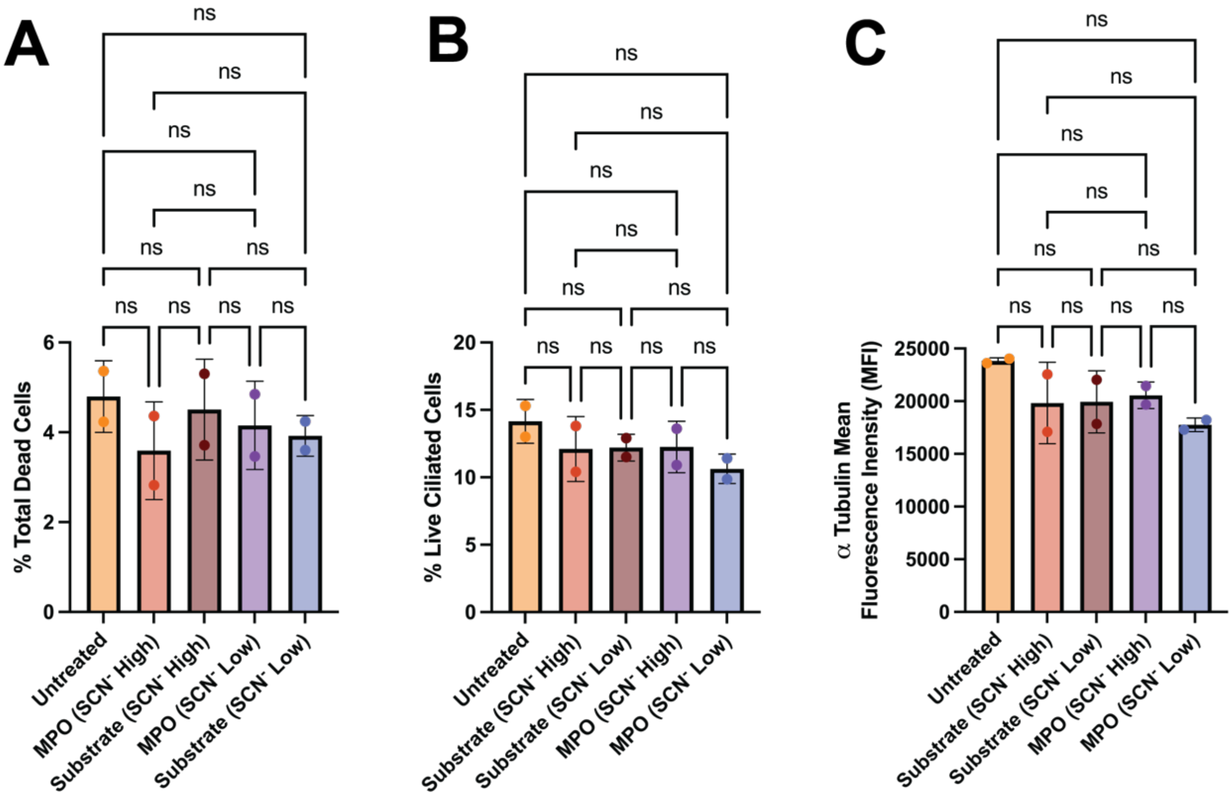


**Figure S1.** Flow cytometry of HAE cells stained to evaluate cell both viability and ciliation using Zombie NIR dye and a fluorophore-conjugated antibody against alpha tubulin, respectively. All samples were initially gated to analyze only the singlet cell population. A) The percentage of all dead cells found in each HAE cell population (Zombie NIR positive). B) The percentage of live ciliated cells in each HAE cell population (Zombie NIR negative, alpha-tubulin positive). C) The ciliary expression of the live ciliated HAE cell population was evaluated using the mean fluorescence intensity. Experimental groups in panels A, B and C were compared statistically using one way ANOVAs and Tukey's multiple comparison tests (ns, no significance).

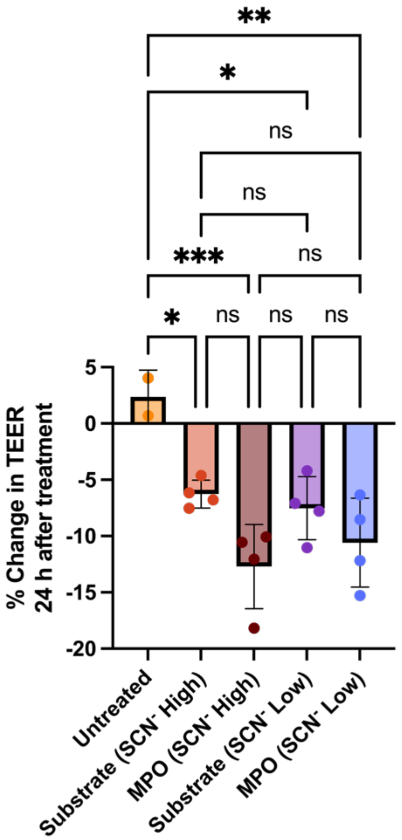


**Figure S2.** Measurements of TEER were performed on each HAE culture pre-treatment and 24 hours following treatment with substrate-only or MPO-substrate solutions. The percent change in TEER values for each individual HAE culture before and after treatment was calculated and plotted. Statistical significance was determined using a one-way ANOVA with Tukey's multiple comparisons test (ns, no significance, * $p < 0.05$, ** $p < 0.01$, *** $p < 0.001$).

.

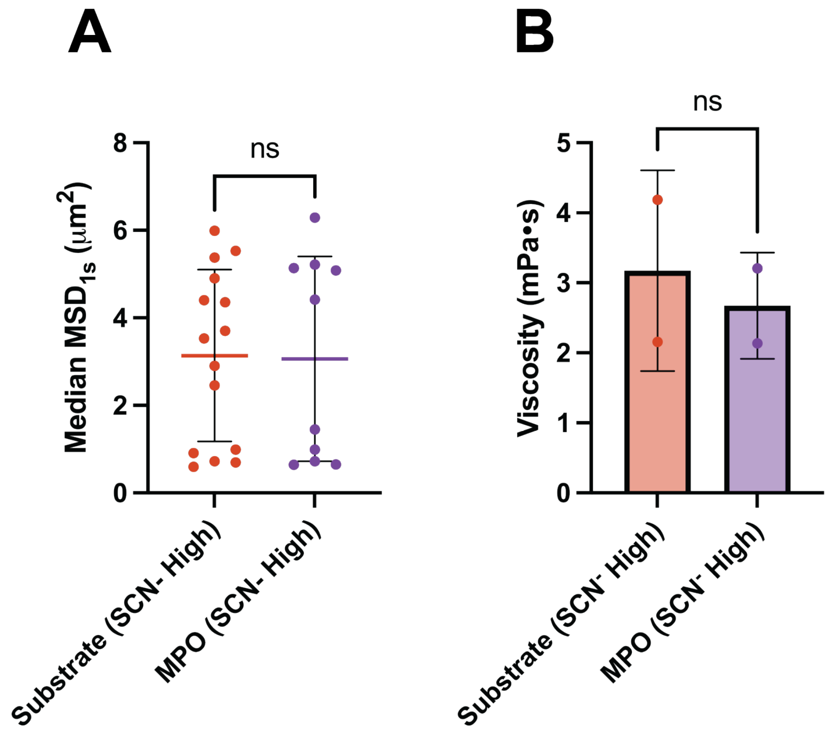


**Figure S3. Microrheology and bulk viscosity of mucus after exogenous treatment with MPO in the absence of cells.** Mucus was harvested from BCi-NS1.1 cultures by washing with PBS and isolating the >100 kDa fraction using centrifugal filtration. The mucus was treated with 30 μg/mL MPO dispersed in buffer at 'SCN⁻ high' reaction condition. **(A)** Results of particle tracking microrheology in substrate-only and MPO-substrate conditions (n = 3 replicates per condition; 5 randomly selected regions of each individual gel were imaged). Data reported as median mean squared displacement (MSD) at 1 second (median $MSD_{1s}$) using PEGylated 100 nm nanoparticles as probes. **(B)** Viscosity of the whole apical wash samples from each treatment group quantified using a microviscometer.

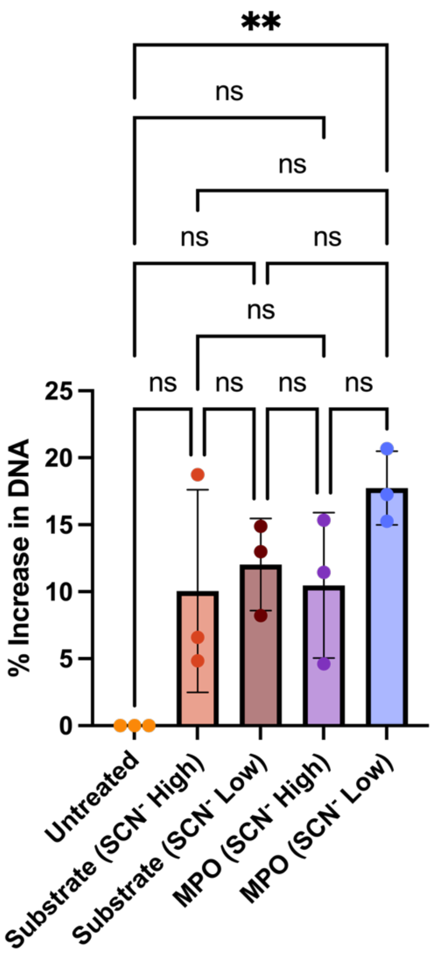


**Figure S4.** An alternate method of DNA quantification in the whole apical wash was used in which Sytox green fluorescent DNA stain was added to the samples. The percent change in fluorescence values was calculated and plotted. Statistical significance was determined using a one-way ANOVA with Tukey's multiple comparisons test (ns, no significance, ** $p < 0.01$).

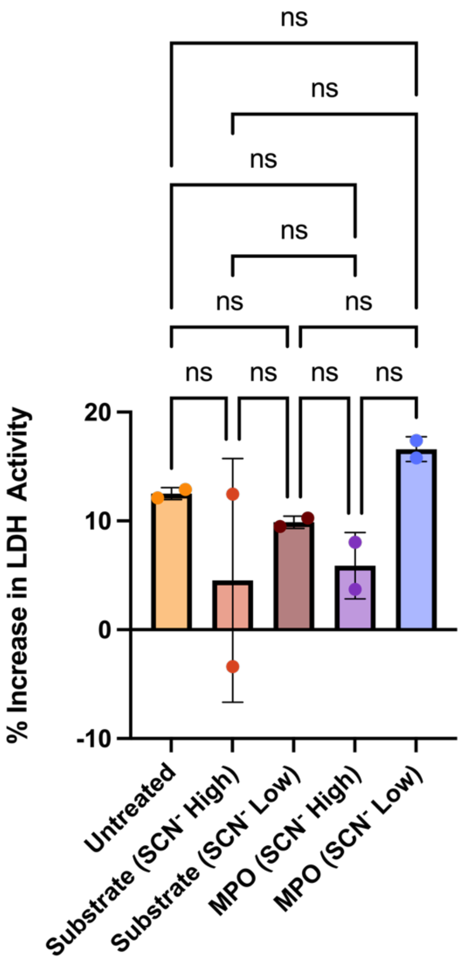


**Figure S5.** The LDH activity in the basolateral media of each individual HAE culture was quantified using a colorimetric assay kit both 24 hours prior to treatment and 1 hour after treatment with substrate-only or MPO-substrate solutions. The percent change in the absorbance values was calculated and plotted. A one-way ANOVA with Tukey's multiple comparisons test was performed as the statistical analysis (ns, no significance).

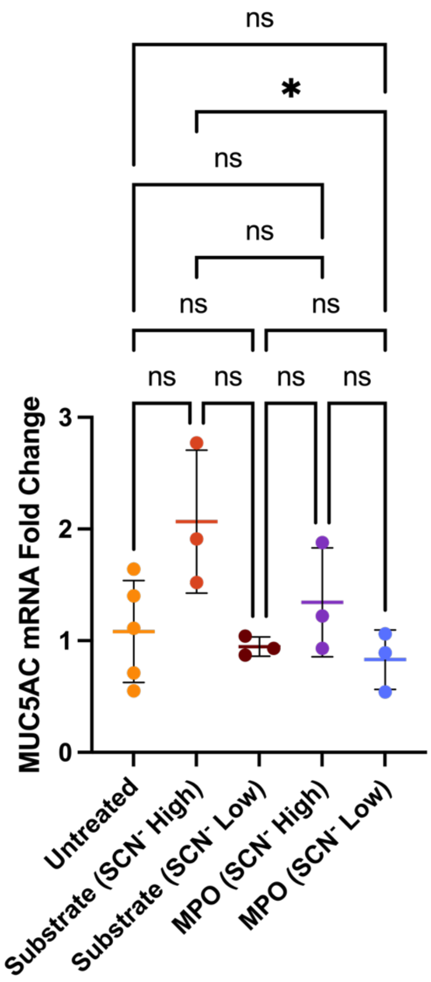


**Figure S6.** The fold change in MUC5AC mRNA expression was quantified using qPCR following 24 hours of treatment with substrate-only or MPO-substrate solutions. The statistical analysis performed was a one-way ANOVA with Tukey's multiple comparisons test (ns, no significance, * $p < 0.05$).
